# Atox1-Cyclin D1 Loop Activity is Critical for Survival of Tumor Cells with Inactivated *TP53*

**DOI:** 10.1101/2024.02.14.580254

**Authors:** O.A. Kuchur, S.S. Pogodaeva, A.V. Shcherbakova, S.A. Tsymbal

## Abstract

The search for relevant molecular targets is one of the main tasks of modern tumor chemotherapy. To successfully achieve this, it is necessary to have the most complete understanding of the functioning of the cell’s transcriptional apparatus, particularly related to proliferation. The p53 protein plays an important role in regulating processes such as apoptosis, repair, and cell division, and the loss of its functionality often accompanies various types of tumors and contributes to the development of chemoresistance. Additionally, the proliferative activity of tumor cells is closely related to the metabolism of transition metals. For example, the metallochaperone Atox1 – a copper transporter protein - acts as a transcription activator for cyclin D1, promoting progression through the G_1_/S phase of the cell cycle. On the other hand, p53 suppresses cyclin D1 at the transcriptional level, thereby these proteins have divergent effects on cell cycle progression. However, the contribution of the interaction between these proteins to cell survival is poorly understood. This work demonstrates that there not only exists a positive feedback loop between Atox1 and cyclin D1, but also that the activity of this loop depends on the status of the *TP53* gene. Upon inactivation of *TP53* in A549 and HepG2 cell lines, the expression of *ATOX1* and *CCND1* genes is enhanced, and their suppression in these cells leads to pronounced apoptosis. This fundamental observation may be useful in selecting more precise interventions for combined therapy of p53-negative tumors.

## INTRODUCTION

The search and identification of specific genetic changes characteristic of tumor cells provides the opportunity to select more precise effects on them to reduce negative side effects while maintaining the same effectiveness of therapy. In this regard, proteins and factors associated with the process of cell division and regulation of individual phases of the cell cycle are especially interesting, since accelerated proliferation is one of the striking distinguishing features of tumor cells [1–3]. One of the key tumor suppressors that modulates the activity of dozens of cascades responsible for normal cell development, apoptotic and repair processes is the p53 protein [4, 5]. This also leads to the problem - tissue malignancy is often accompanied by the acquisition of p53 driver mutations, as a result of which tumor suppressor functions are lost, which entails uncontrolled division, weakening of repair processes, and the formation of chemo- or radioresistance, for example, in breast and lung tumors [6–8]. Changes in p53 functions can be caused not only by mutations in the *TP53* gene, but also epigenetically (phosphorylation by Cdc2, JNK1, protein kinase C) [9], as well as mutations of the *TP53* regulators - TP53BP1, Chek2, ATM, Usp28, which leads to a weakening of the control of cellular division [10]. Approximately 50% of tumors have been shown to have missense mutations, point mutations, *TP53* deletions, or splicing abnormalities. In most cases, *TP53* mutations occur in exons 5-8, which encode the DNA-binding domain of the protein. This explains the pro-oncogenic function more than 80% of p53 missense mutations [11–14].

In addition to participating in the processes of repair and apoptosis, one of the main functions of p53 is the regulation of the *CDKN1A* gene, which encodes the inhibitor of cyclin-dependent kinase 1A, the p21 protein. In cooperation with p53, this protein suppresses the activity of the CCNA1/CCNB1-CDK1/CDK2 and CCND1-CDK4/6 complexes, which causes cell cycle arrest at the G_1_/S boundary and in the G_2_/M phase, promoting DNA repair. Inactivation of p53 leads to overexpression of the *CCND1* gene encoding cyclin D1, loss of control of cell cycle phase changes, and accumulation of damage [15, 16]. Therefore, cyclin D1 in complexes with CDK4 and -6 is an important therapeutic target [17, 18].

Along with the regulation of the cell cycle and responses to DNA damage, p53 is a modulator of metabolic cascades. The role of p53 in controlling the expression of genes for glucose metabolism [19, 20], respiration [21], and metabolism of transition metals, especially copper [22–24], is known. The connection of p53 with copper metabolism is also important due to the discovery of the role of this metal in tumor biology [25–28]. A new type of cell death has been discovered – cuproptosis, which is associated with p53-dependent cascades [29]. The distribution and transport of copper ions in cells is regulated by an 8-9 kDa metallochaperone Atox1 (Antioxidant protein 1) [30]. After the copper transporter Ctr1 transfers it into the cell, Atox1-mediated localization of Cu to the ATPases ATP7A and ATP7B occurs in the trans-Golgi network for inclusion in copper-dependent enzymes: superoxide dismutase SOD1, lipoxygenase LOX, etc. [31, 32].

One of the nontrivial features of Atox1 is the participation of a metallochaperone in the transcriptional regulation of the cyclin D1 gene, which directly links copper metabolism with proliferation. Thus, the addition of copper complexes to the medium of mouse embryonic fibroblasts (MEFs) increased cyclin D1 mRNA; in cells with knockout of both Atox1 alleles, this effect was absent. Inactivation of Atox1 resulted in suppression of proliferation, decreased cyclin D1 levels, and arrest of the transition from G_1_ to S. Analysis of gene regulatory regions showed that copper stimulates the binding of Atox1 to a previously undescribed cis-element in the *CCND1* promoter [33, 34].

The regulation of Atox1 and the intensity of copper ions transportation into the nucleus depend on p53: in MEF cells with knockdown of the *TP53* gene, the functions of Atox1 and Atox1-mediated regulation of *CCND1* are reduced [35, 36]. On the other hand, our previous study shows that in cells of epithelial origin, inactivation (knockout or knockdown) of *TP53* induces *ATOX1* and increases the amount of its product [37], which may indicate tissue-dependent regulation.

Participation in the regulation of the most important processes in the cell - the balance of oxidation-reduction and changes in phases of the cell cycle – suggests the presence of a functional connection between p53, Atox1 and *CCND1*/cyclin D1. Each mechanism individually and their combinations are important for achieving the main goal of drug therapy – irreversible death of as much of the tumor cell population as possible.

The purpose of this work is to establish the functional relationships of *TP53, ATOX1* and *CCND1*/cyclin D1 in the responses of human tumor cells to “chemotherapeutic stress”: antitumor compounds. Bleomycin is an inducer of single- and double-strand DNA breaks, as well as reactive oxygen species [38, 39]. Both processes cause activation of p53-dependent responses. Palbociclib, a specific inhibitor of CDK4/6, was selected for the functional inhibition of cyclin D1, a protein partner of these protein kinases [40]. How are combinations of these effects realized in cells with intact and nonfunctional p53? Is it possible to discover a combination of these factors in which cell death is especially effective?

## MATERIALS AND METHODS

### Cell lines and culture conditions

The lines of wild-type lung adenocarcinoma A549 (Biolot, Russia) and hepatocellular carcinoma HepG2 (Russian collection of cell cultures of the Institute of Cytology of the Russian Academy of Sciences) were used. Subline A549TP53^-/-^ (*TP53* inactivated by the CRISPR-Cas9 method) was obtained at the Institute of Gene Biology, Russian Academy of Sciences, Ph.D. Bruter A.V [41]. Cells were cultured in Dulbecco’s modified Eagle medium - DMEM (Biolot) supplemented with 10% fetal bovine serum (PAA, USA), 2 mM L-glutamine and 100 U/ml gentamicin (Biolot). Cells in the logarithmic growth phase were used in the experiments. 6-well plates (BioFil, China; 10^5^ cells per well) were used during the experiments.

### Pharmacological effects and IC50 determination for tumor cell lines

To determine the degree of cytotoxicity, the MTT test was used; the effect was determined by the enzymatic activity of mitochondrial enzymes [42]. Briefly, cells were plated into a 96-well plate at a concentration of 5 thousand cells/well, after which a serial dilution was carried out: bleomycin (Bleo) in concentrations from 7-0.012 mM and palbociclib (Palbo) in concentrations from 25 mM-0.098 mM were added to the cells after 24 hours. The cells were incubated with the drugs for 72 hours, after which MTT (Thermo FS, USA) reagent was added to them for 2 hours, then the medium was taken and DMSO (PanEco, Russia) was added to dissolve the formazan crystals formed in the mitochondria. For qPCR analysis and immunoblotting, palbociclib and bleomycin were added for 24 hours, in the case of cell cycle analysis for 48 hours.

### Characteristics of the combined drugs effect

Based on the results obtained during the MTT analysis, the characteristics of possible combination effects of the drugs were assessed. For this purpose, drug combination indices (CI) were calculated using the original mathematical apparatus proposed by Chou [43]. In this way, the synergistic, additive and antagonistic interaction of drugs in relation to cells was determined.

The index was calculated using the formula:

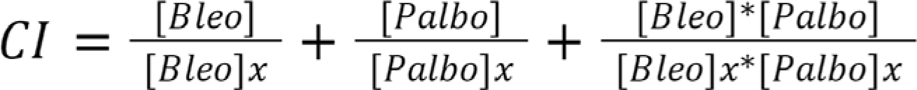

where [Bleo] and [Palbo] – the selected doses of bleomycin and palbociclib, respectively. [Bleo]x and [Palbo]x - IC50 of bleomycin and palbociclib. According to the methodology used for calculating the index, with a value of CI < 0.9, a synergistic interaction of drugs is observed, with 0.9 < CI < 1.1 - additive, with CI > 1.1 - antagonistic.

### siRNA transfection

GenJect reagent (Molecta, Russia) was used for transfection of siRNA (125 nmol) according to the manufacturer’s instructions in DMEM medium without the addition of serum and glutamine. Transfection was carried out 24 hours before treatment with drugs, the sequences are indicated in Table 1. Cell density was 200 thousand cells per well; before adding drugs, the medium was replaced with growth medium. The green fluorescent protein (GFP) mRNA sequence was used as a gene-independent control. All oligonucleotide sequences were ordered from DNA-Synthesis, Russia.

**Table 1.**
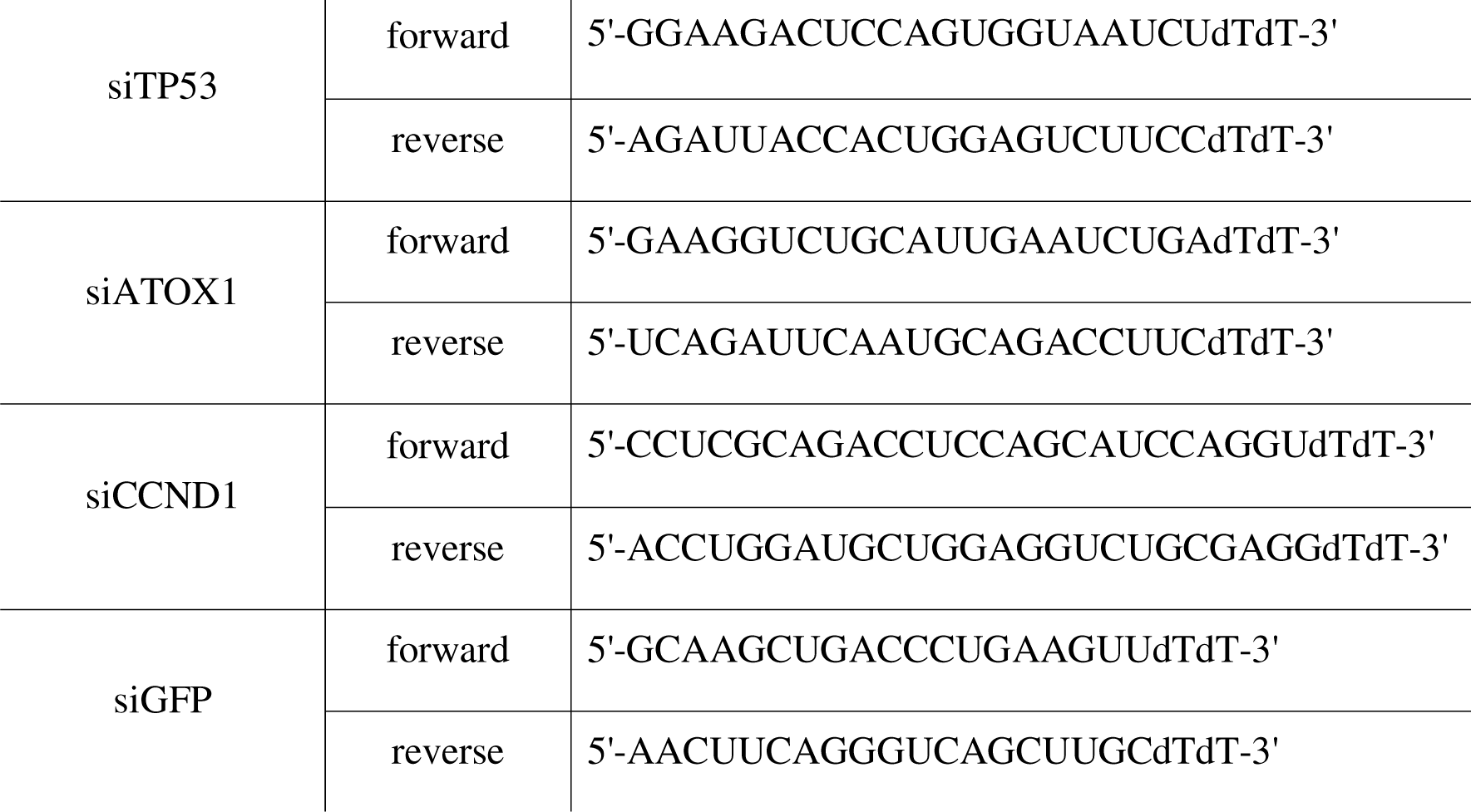
siRNA sequences.

### Cell cycle analysis

The distribution of cell cycle phases was observed using a CytoFlex B2-R2-V0 flow cytometer (Beckman Coulter, USA) after fixation in 80% ice-cold ethanol. Cells were pelleted by centrifugation and lysed in a buffer containing 25 μg/ml propidium iodide and 100 μg/ml RNase (Thermo FS, USA). The recording was carried out in the TRITC channel, 20,000 events were accumulated for each sample.

### Characterization of cell death

To determine the apoptotic or necrotic nature of cell death, cells were double stained using oxazolium yellow (YO-PRO™-1) and propidium iodide (PI). Drugs and siRNA were added according to the previously described experimental plan. After transfection and incubation with drugs, the cells were detached from the plate, centrifuged for 1 minute at 500g, the sediment was dissolved in a warm medium, 10 μM YO-PRO™-1 was added to the samples, and incubated for 30 minutes at room temperature. Afterwards, 4 µM PI was added to the samples and incubated for 5-10 minutes. The results were recorded using a CytoFlex B2-R2-V0 flow cytometer (Beckman Coulter, USA) in the FITC and TRITC channels, respectively.

### RNA extraction, reverse transcription and real-time PCR

Total RNA was extracted using ExtractRNA buffer (Evrogen, Russia) according to the manufacturer’s protocol. For reverse transcription, 2 μg of total RNA and a reaction mixture with MMLV reverse transcriptase (Evrogen, Russia) were used. The analysis was carried out using a CFX96 Touch thermal cycler (Bio-rad, USA). Primer sequences for qPCR are listed in Table 2. Comparison of gene expression in qPCR was determined by ΔCt, where Ct (threshold cycle) is the cycle at which the fluorescence level reaches a certain value (selected threshold), and Δ is the change in the cDNA of the gene under study relative to the *HPRT* gene (hypoxanthine-guanine phosphoribosyltransferase, normalization). In all groups, differences with the control were significant at p ≤ 0.01 (one-way ANOVA test).

**Table 2.**
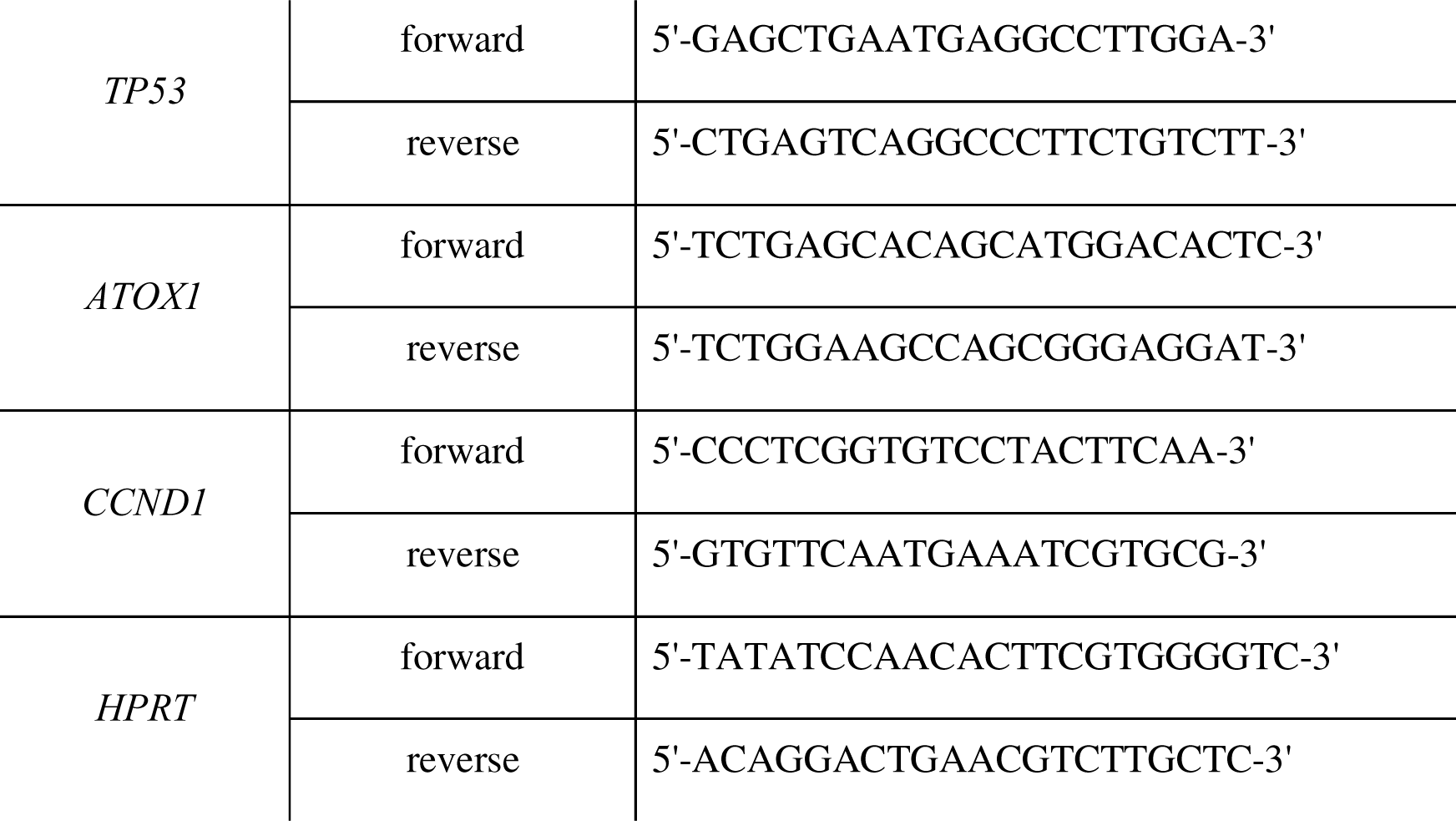
Primer sequences.

### PAGE and immunoblotting

Cells were lysed in RIPA buffer (50 mM TrisHCl (pH 8.0), 150 mM NaCl, 1% NP40, 0.1% SDS) supplemented with a protease inhibitor mixture (Sigma-Aldrich, USA). Protein concentration was determined using Bradford reagent (Sigma-Aldrich, USA) on a spectrophotometer (Infinite F50 Tecan, Switzerland) at wavelengths of 595 nm and 450 nm [44]. Protein electrophoresis was performed using a 10-12% polyacrylamide gel containing 10% sodium dodecyl sulfate (30 μg of total protein per track). After electrophoresis, the proteins were transferred to a nitrocellulose membrane (Amersham, USA) and incubated overnight at 4°C with primary antibodies to p53, cyclin D and Atox1 (dilution 1:500–1:2000 in TBST). Antibodies to β-actin (1:1000) were used to control the amount of protein added. The membrane was washed from primary antibodies and mouse or rabbit IgG secondary antibodies conjugated with horseradish peroxidase (1:1000) were added. All antibodies were purchased from Cell Signaling Tech., USA. Chemiluminescent signals were recorded using the ChemiDoc Touch gel documentation system (Bio-Rad, USA). Densitometric analysis to estimate the relative abundance of proteins was performed using ImageJ (Plot Lanes tool).

### Statistical analysis

The GraphPad Prism 8.0 program (GraphPad Software, USA) was used to construct graphs and process the results. The results were processed using one-way ANOVA analysis; differences were statistically significant at *p ≤ 0.05, **p ≤ 0.01. The number of replicates for all experiments was n = 3.

## RESULTS

In preliminary experiments, the viability of cell cultures was assessed using the MTT assay upon treatment with bleomycin and palbociclib. The half-maximal inhibitory concentrations (IC50) were obtained as follows: for bleomycin in A549 cells, the IC50 value was 17.32 μM; in A549TP53^-/-^, the IC50 was lower at 3.61 μM. This difference is presumably associated with the protective functions of the tumor suppressor p53, which activates DNA repair processes upon damage. For palbociclib in A549 cells the IC50 was 8.84 μM (Fig. 1A), while in A549TP53^-/-^ cells it was 3.5 μM (Fig. 1B). In HepG2 cells, the IC50 values were 1.59 μM for bleomycin and 1.2 μM for palbociclib (Fig. 1C).

**Fig. 1.**
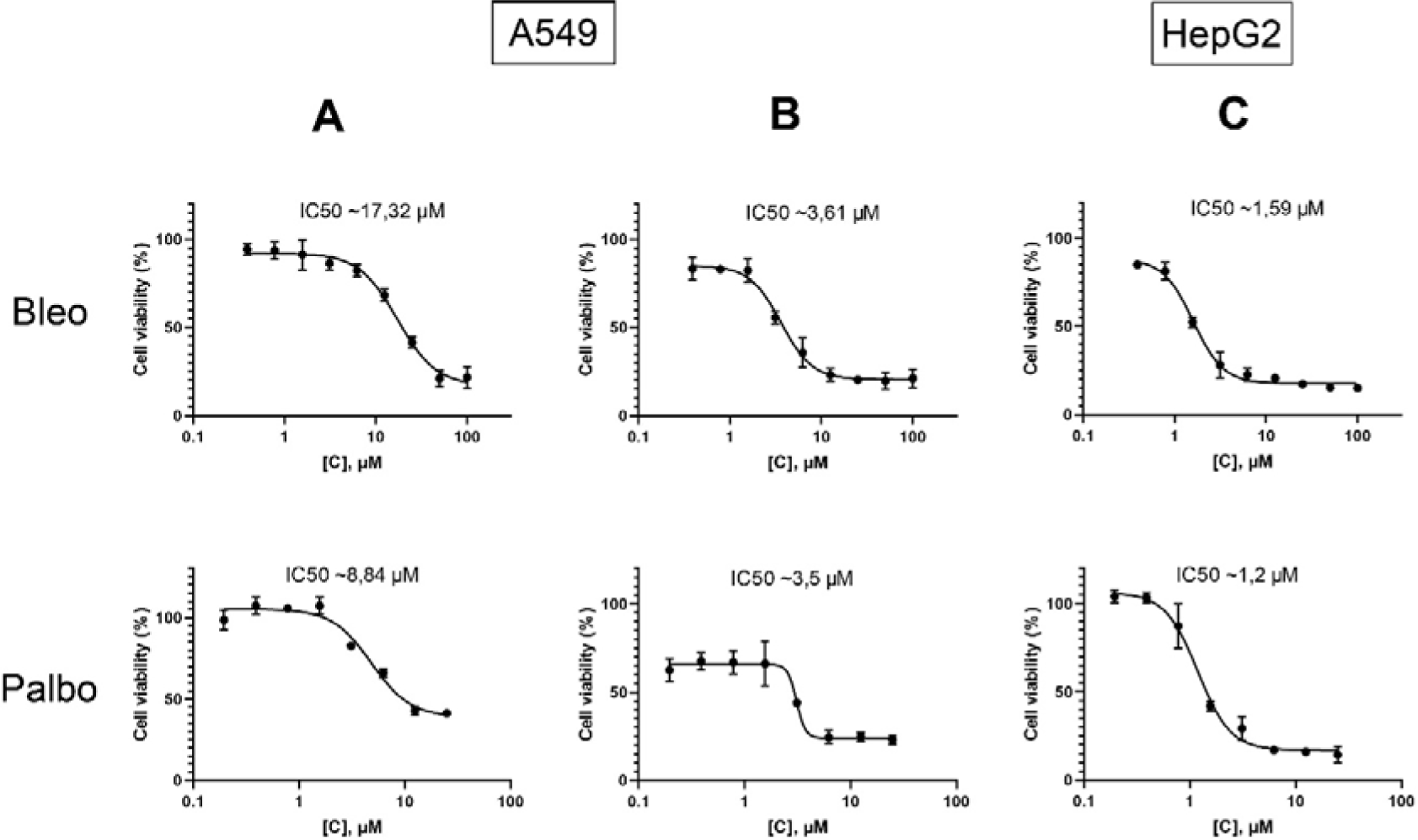
Results of the MTT assay to determine the IC50 of bleomycin (Bleo) and palbociclib (Palbo) drugs, 72 hours after drug addition: A - titration of drugs for the A549 cell line. B - titration of drugs for the A549 TP53^-/-^ cell line. C - titration of drugs for the HepG2 cell line. Cells without treatment were used as controls. Control was considered as 100% viability. The mean values from 3 independent measurements are shown ± SEM.

### The combined exposure of bleomycin and palbociclib exhibits a synergistic effect

Characterization of the combinatorial exposure of drugs allows for the investigation of interactions between two or more substances that were simultaneously or sequentially introduced to cell cultures. The following concentrations of simultaneously added drugs to the cells were selected for the study based on the obtained IC50 values for each cell line: for A549, the concentration of bleomycin was 5 μM, and the concentration of palbociclib was 2.5 μM. The calculated combination index (CI) for this combination of bleomycin and palbociclib is approximately 0.641. In the case of the A549TP53^-/-^ cell line, values of 1.5 μM and 1 μM were chosen for bleomycin and palbociclib, respectively. The CI in this case is 0.813. In both instances, the obtained values indicate a synergistic effect of the combination according to the formula provided in the Materials and Methods section.

For the HepG2 cell line, the following concentrations of substances were selected based on the obtained IC50 values and preliminary experiments: the dose of bleomycin was 0.5 μM, and the dose of palbociclib was also 0.5 μM. At these concentrations, the combination index (CI) is approximately 0.861, which also suggests a synergistic effect of the two drugs.

### Bleomycin-induced p53 activation leads to the suppression of Atox1 and cyclin D1, while the action of palbociclib suppresses Atox1

To test the hypothesis of the interplay between *ATOX1* and *CCND1*, both under the control of *TP53* at the transcriptional level, qPCR analysis was conducted on the respective genes (Fig. 2A). Treatment of A549 cells with bleomycin increased *TP53* expression by 4-5 times and simultaneously reduced *ATOX1* and *CCND1* mRNA levels by half. Addition of palbociclib did not alter *TP53* and *CCND1* expression, while *ATOX1* mRNA decreased by an average of 2.5 times. Comparable results were obtained in the HepG2 cell line. The combination of palbociclib and bleomycin activated *TP53* by 7-fold in A549 and 3.5-fold in HepG2 compared to untreated control, leading to decreased expression of *ATOX1* and *CCND1*.

**Fig. 2.**
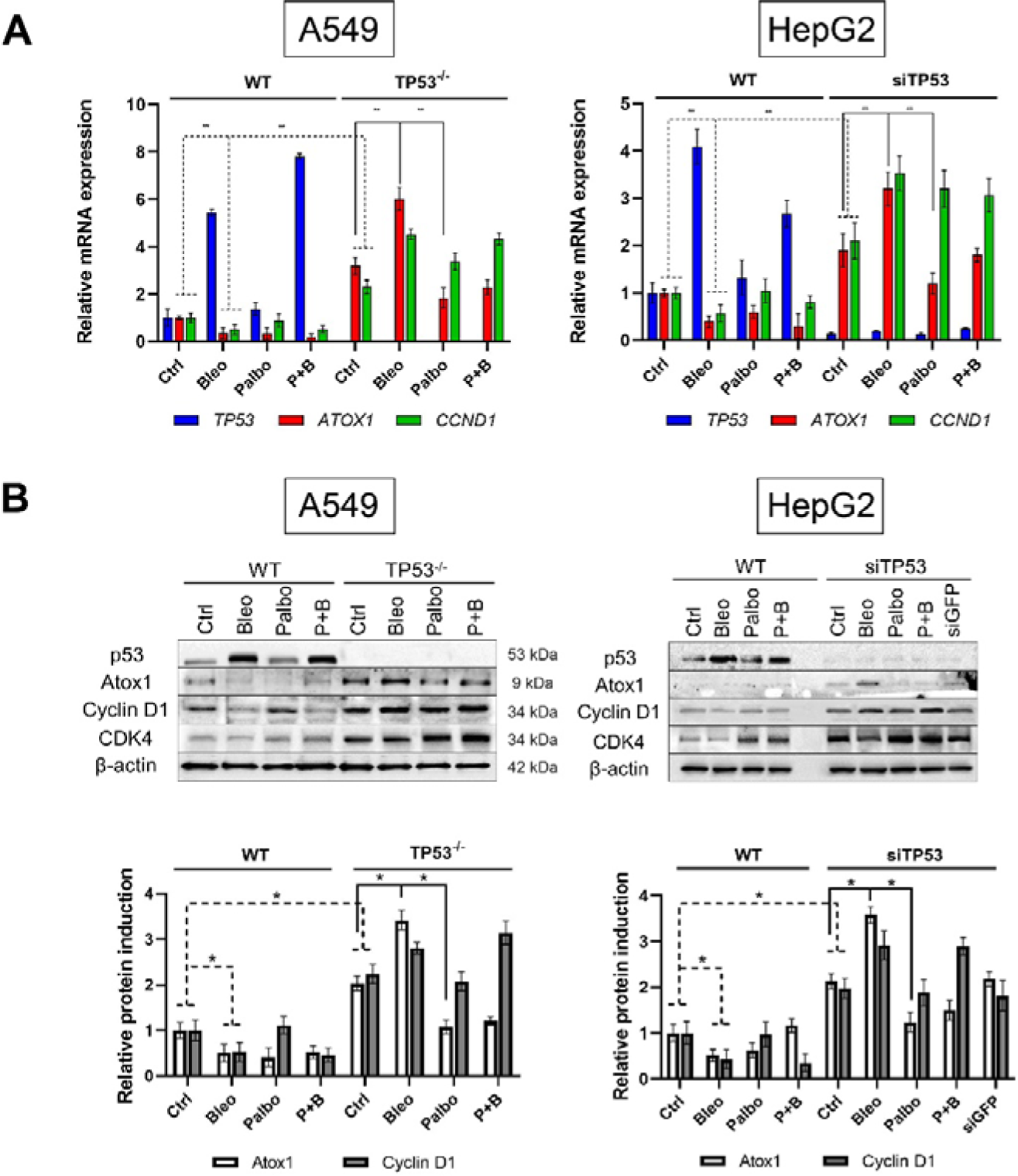
A - gene expression of *TP53, ATOX1*, and *CCND1* in A549 (WT and TP53^-/-^) and HepG2 (WT and siTP53) cell lines after 24 hours of treatment with palbociclib (Palbo), bleomycin (Bleo) at their IC50 concentrations, and also combination of exposure (P+B). Normalization to *HPRT* gene expression levels. *GFP*-specific siRNA was used as a gene-specific control. The intact control was set as 1.0 for all genes. Mean values of three independent measurements ± SEM are shown. ** - Significant differences between the indicated values (p-value < 0.01, one-way ANOVA). B - induction of p53, Atox1, cyclin D1, and CDK4 proteins in A549 (WT and TP53^-/-^) and HepG2 (WT and siTP53) cells after 24 hours of treatment with Palbo (IC50), Bleo (IC50), and their combination (P+B), with β-actin as loading control. Corresponding densitometric analysis graphs are presented below. Mean values for each sample ± SEM are indicated. * - Significant differences between the indicated values (p < 0.05).

In preliminary experiments by Khamidullina A.I. et al., the generation and validation of a *TP53* knockout line in A549 using the CRISPR/Cas9 system were described, with selection based on resistance to nutlin-3 [41]. The knockout phenotype was confirmed by the complete absence of the p53 protein via qPCR and immunoblotting, reduced expression of the p53-dependent gene *CDKN1A*, and altered sensitivity to DNA-damaging antitumor drugs. In our study, in *TP53*-inactivated cell lines (either stable: deletion of both alleles in the A549TP53^-/-^ subline, or transient: anti-sense RNAs in the HepG2 line), basal expression of *ATOX1* and *CCND1* was increased 2-3 times compared to corresponding cells with intact *TP53*. Treatment with bleomycin resulted in effects opposite to those observed in cells with intact p53: levels of both *ATOX1* and *CCND1* expression increased by 50%. Palbociclib caused a 2-fold decrease in *ATOX1* mRNA. This same compound, in combination with bleomycin, also reduced *ATOX1* mRNA, counteracting the activating effect of bleomycin. All three treatments (bleomycin, palbociclib, and their combination) increased *CCND1* expression relative to untreated cells with non-functional p53 (Fig. 2A). Thus, palbociclib, as a disruptor of the CCND1-CDK4/6 complex, reduces *ATOX1* expression, suggesting a role for cyclin D1 as a direct or indirect transcriptional regulator of this gene. Importantly, this relationship varies depending on the presence of functional p53. The regulatory axis *ATOX1-CCND1-TP53* is identified for the first time.

The assessment of protein induction levels confirmed the qPCR results (Fig. 2B). In cell lines with intact p53, bleomycin led to the accumulation of p53 while decreasing Atox1 and cyclin D1. Treatment with palbociclib reduced the level of Atox1 by 2-3 times without affecting the levels of p53 and cyclin D1, possibly because the drug does not interfere with the synthesis processes of cyclin D1, only affecting its function. In cells with inactivated *TP53*, the relative amounts of Atox1 and cyclin D1 increased by 2-fold, with bleomycin further increasing Atox1 and cyclin D1. Palbociclib significantly reduced Atox1 without affecting cyclin D1. Similar to the mRNA level, the combination of bleomycin and palbociclib did not alter the levels of Atox1 and cyclin D1 proteins compared to single-agent bleomycin treatment. Additionally, an analysis of CDK4 induction, a molecular partner of cyclin D1 in the CCND1-CDK4/6 complex, was conducted. In cell lines with inactivated TP53, the level of CDK4 was approximately 2-2.5 times higher than in wild-type cells, as previously observed [45]. Upon addition of bleomycin, CDK4 behaved differently from Atox1 and cyclin D1 - its induction was slightly reduced, but when combined with drugs, its level remained higher than the intact control. However, these effects were not pronounced and did not exceed 2-fold deviations from untreated cells. The results for A549 and HepG2 cell lines were similar for all proteins (Fig. 2B).

Based on the results presented above, we can conclude that the expression of *ATOX1* and *CCND1* is dependent on the p53 status, as its presence inhibits both genes. It is noteworthy that the response to genotoxic stress is highlighted: the addition of the cytostatic bleomycin suppresses the expression of *ATOX1* and *CCND1* in wild-type cells and enhances it when *TP53* is inactive. There may exist a mechanism of coregulation between cyclin D1 and Atox1: while it has been previously shown that Atox1 can act as a transcription factor for cyclin D1, here we observe a decrease in *ATOX1* expression when the function of cyclin D1 is suppressed, indicating a potential positive feedback loop between these proteins.

### Interdependent regulation of *ATOX1* and *CCND1* activity

We investigated the possibility of coregulating the genes *ATOX1* and *CCND1* using specific inhibition of mRNA of each gene by small interfering RNA (siRNA). As shown in Fig. 3A, in cells with intact p53, 24 hours after transfection with the corresponding siRNAs, a reduction in *ATOX1* and *CCND1* transcripts to 90-95% of control levels was observed (transfection reagent (GenJect) without siRNA or introduction of control RNA to siGFP). Simultaneous or separate transfection of siRNA targeting *ATOX1* and *CCND1* did not affect the level of *TP53* mRNA. Importantly, siCCND1 led to a significant - 4-5 fold - decrease in *ATOX1* mRNA, while adding siATOX1 reduced *CCND1* expression. Bleomycin increased *TP53* mRNA and decreased *ATOX1* and *CCND1* transcripts to levels similar to the effects of siRNA targeting *ATOX1* and *CCND1*. The results were comparable in both cell lines.

**Fig. 3.**
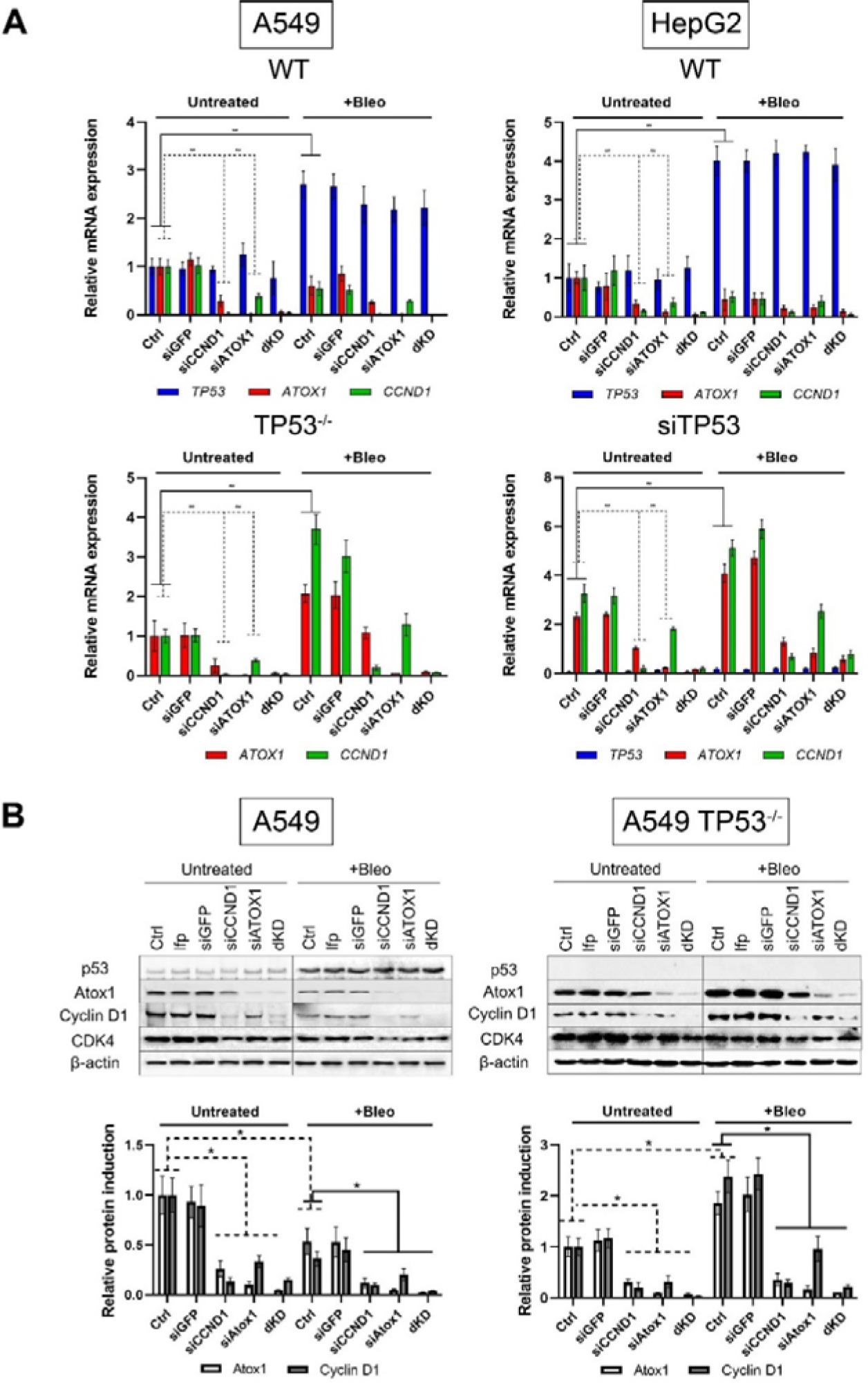
A – Expression of *TP53, ATOX1*, and *CCND1* genes in A549 (WT and TP53^-/-^) and HepG2 (WT and siTP53) cell lines following siRNA-mediated suppression of *CCND1* (siCCND1), *ATOX1* (siATOX1), and their combined inhibition (dKD), 24 hours after treatment with bleomycin (Bleo, IC50 for each subline, respectively). Normalization to *HPRT* gene expression level. *GFP* siRNA was used as a gene-specific control. Intact control set at 1.0 for all genes. Mean values from 3 independent measurements ± SEM are shown. ** – Significant differences between the specified values (p-value < 0.01, one-way ANOVA). B – Induction of p53, Atox1, cyclin D1, and CDK4 proteins in A549 (WT and TP53^-/-^) and HepG2 (WT and siTP53) cells, 24 hours after treatment with bleomycin (Bleo, IC50). β-actin was used as loading control. Corresponding densitometric analysis graphs are presented below. Mean values for each sample ± SEM are shown. * – Significant differences between the specified values (p-value < 0.05, one-way ANOVA).

*TP53* inactivation leads to a more pronounced interplay of the investigated mechanisms. Suppression of *CCND1* reduced *ATOX1* expression by 3-4 fold in both cell lines. siATOX1 caused a > 2.5-fold decrease in *CCND1* expression. Addition of bleomycin activated both *ATOX1* and *CCND1*, but did not negate the mutual regulation of these genes (Fig. 3A). Thus, the mutual influence of these two factors occurs in unstimulated cells and in response to bleomycin.

Protein analysis confirms the observations obtained in qPCR: a decrease in *ATOX1* mRNA when *CCND1* is suppressed and vice versa. In cells with intact p53, the simultaneous suppression of *ATOX1* and *CCND1* is particularly pronounced: the mRNA levels of each gene are reduced by 10-15 times compared to control values (Fig. 3B). In sublines with nonfunctional p53, suppression of *CCND1* reduced the amount of Atox1 by ∼4-fold, and *ATOX1* siRNA reduced the level of cyclin D1 three-fold. Bleomycin caused an increase in both proteins, while the interaction between Atox1 and cyclin D1 was preserved. CDK4 behaved in a similar way: the protein was noticeably higher in the line with inactivated *TP53*; suppression of *CCND1* or *ATOX1* suppressed its induction, but the effects were not so pronounced. Bleomycin in both sublines reduced CDK4 activity, an effect different from that observed in A549TP53^-/-^ cells for cyclin D1 and Atox1 (Fig. 3B).

The established connection between the three mechanisms allows us to raise the question of the role of the p53-Atox1-cyclin D1 axis in the regulation of the cell cycle and cell viability.

### Suppression of Atox1 and cyclin D1 enhances cell death in cells with non-functional p53

The distribution of cell cycle phases in A549 and HepG2 cells (WT p53) is shown in Fig. 4. In untreated cells, the proportion of hypodiploid cells (DNA fragmentation, spontaneous apoptosis) was <5% (subG_1_), with G_1_ phase at 70-75%, and G_2_/M phase at 20-25%. Palbociclib caused an accumulation of cells in the G_1_ phase by 10-15% compared to controls. Bleomycin increased the percentage of apoptotic cells to 12-15%; simultaneously, an accumulation of cells in G_2_/M (40-50%) was observed. The combination of palbociclib and bleomycin led to the following distribution: 20%, 70%, 10% for subG_1_, G_1_, and G_2_/M phases, respectively. Both drugs were administered for 48 hours.

**Fig. 4.**
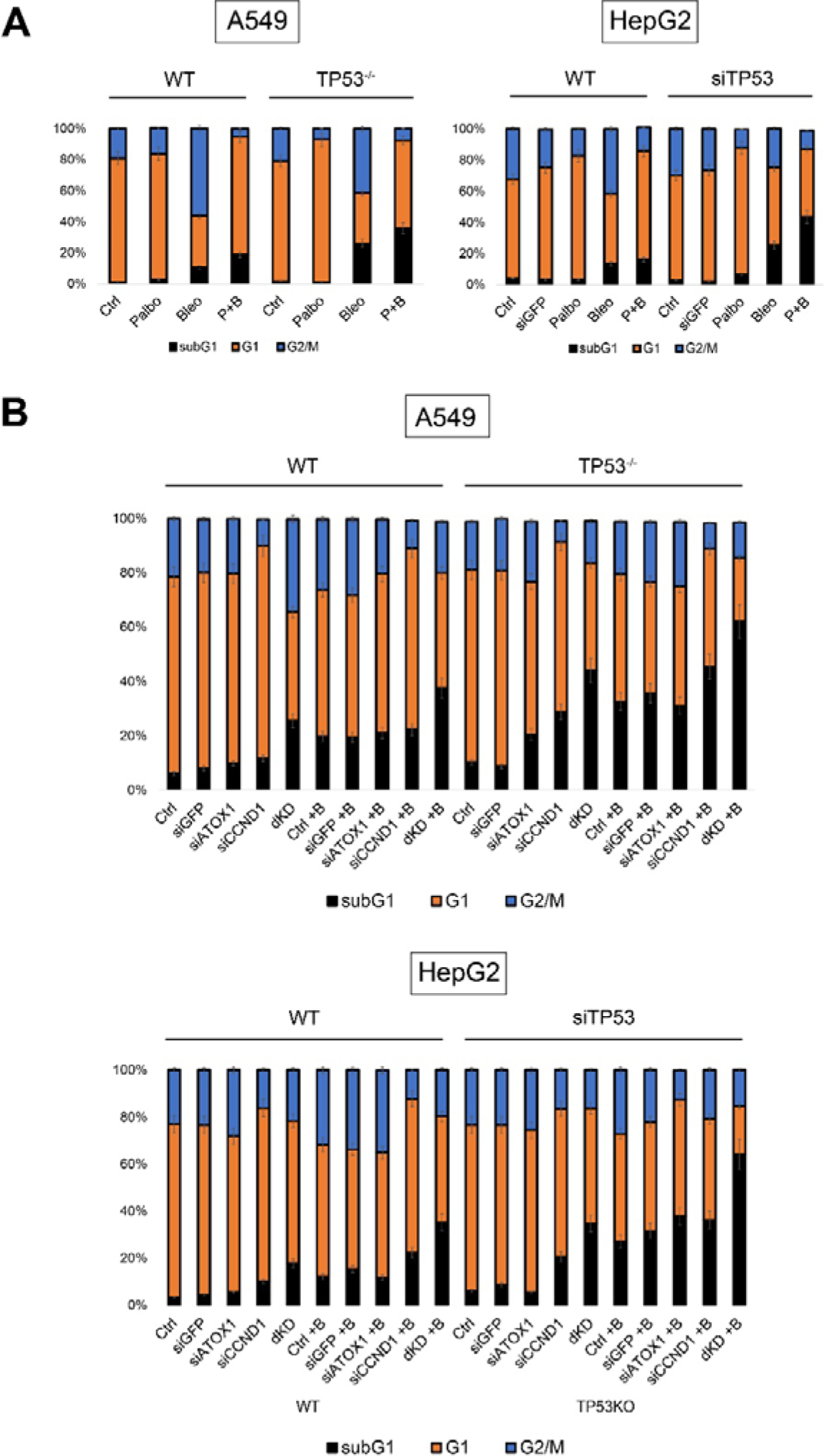
A – Cell cycle analysis upon treatment with Palbo, Bleo, and their combination (P+B) in A549 and HepG2 cell lines with different TP53 statuses, 72 hours after treatment with drugs (IC50 for each subline, respectively). B – Cell viability assay upon sequential siRNA-mediated inactivation (siCCND1, siATOX1), as well as simultaneous suppression of *ATOX1* and *CCND1* (dKD), under intact conditions and upon addition of Bleo (+B) to A549 and HepG2 cell lines. Mean values from 3 independent measurements ± SEM are shown, differences are significant (p < 0.05, one-way ANOVA).

Inactivation of *TP53* did not increase the apoptotic fraction in untreated cells (subG_1_ <3%), palbociclib induced an increase in the G_1_ phase. Bleomycin was more toxic: 25-30% of cells in subG1, 30-40% in G_2_/M. The combination resulted in the following distribution: the subG_1_ fraction almost doubled compared to the corresponding group with intact p53, while the percentage of cells in G_1_ and G_2_/M phases decreased (Fig. 4A). The results were comparable for both cell lines.

Subsequently, siRNA-mediated knockdown of *ATOX1* and *CCND1* individually and in combination (double knockdown, dKD) was performed. In parental cells (intact p53), the distribution of cell cycle phases with single knockdowns of *ATOX1* and *CCND1* did not significantly change; however, simultaneous knockdown (dKD) in both cell lines resulted in 20-25% subG_1_ cells. Addition of bleomycin (48 h) and siRNA targeting *ATOX1* and *CCND1* did not significantly increase the subG_1_ fraction compared to treated samples without inactivation of these genes (20%); however, the combined action of siRNA (dKD) led to the death ∼40% of cells.

In A549TP53^-/-^ subline, the suppression of *ATOX1* and *CCND1* was accompanied by pronounced apoptosis: 25-30% in subG_1_. Double knockdown of *ATOX1* and *CCND1* led to even more significant cell death: 40% in subG_1_, which was comparable to the addition of bleomycin in cells with wild-type *TP53* under double knockdown conditions. Additionally, the proportion of cells in G_2_/M was halved. Genotoxic effects showed similar outcomes: transfection with siATOX1 or siCCND1 followed by bleomycin treatment resulted in the death of 40% of cells. The combination of double knockdown with bleomycin proved lethal for the majority of the cell population: >60% of cells with hypodiploid DNA. These effects were also observed in the HepG2 cell line after *TP53* knockdown (Fig. 4B). Thus, the suppression of *ATOX1* alone does not significantly impact cell viability; however, simultaneous inhibition of *ATOX1* and *CCND1* induces a toxic effect that is enhanced by *TP53* inactivation and the action of antitumor compounds.

### Cell death upon inactivation of ATOX1 and cyclin D1 occurs primarily via apoptosis

Oxazole yellow (YO-PRO-1) is a dye that stains cells undergoing apoptosis; the dye is unable to penetrate inside the cell due to its relatively large molecular weight (630 Da). Apoptotic processes disrupt membrane integrity, allowing YO-PRO-1 to enter the cells, with its fluorescence detected in the green spectrum. Propidium iodide (PI), on the other hand, serves as a marker for necrotic cells by binding to nucleic acids, particularly DNA, which is released from the nucleus. PI fluorescence is detected in the red spectrum.

Continuing from the previous section, cell death of A549 and HepG2 cells was characterized after the suppression of *TP53*, *ATOX1* and *CCND1* genes in the presence/absence of bleomycin. As a control for necrotic cell death, cells were incubated at 65°C for 10 minutes (not shown on the graph). Figure 5 shows the distribution of cell populations based on the nature of cell death. While controls and samples with inactivation of a single gene (*ATOX1* or *CCND1*) did not exhibit significant cell death, clear apoptotic/late apoptotic cell death was observed for wild-type A549 cells treated with bleomycin under simultaneous inactivation of *ATOX1* and *CCND1* (dKD+B), 40% fraction. The A549 cell subline with a non-functional *TP53* gene was more sensitive to bleomycin: apoptotic cell death was evident in controls (38%) and upon addition of siATOX1 (35%) or siCCND1 (40%). Suppression of *TP53*, *ATOX1*, *CCND1* genes in combination with bleomycin leads to the death of 60% of cells, with approximately 15% exhibiting necrotic characteristics.

**Figure 5.**
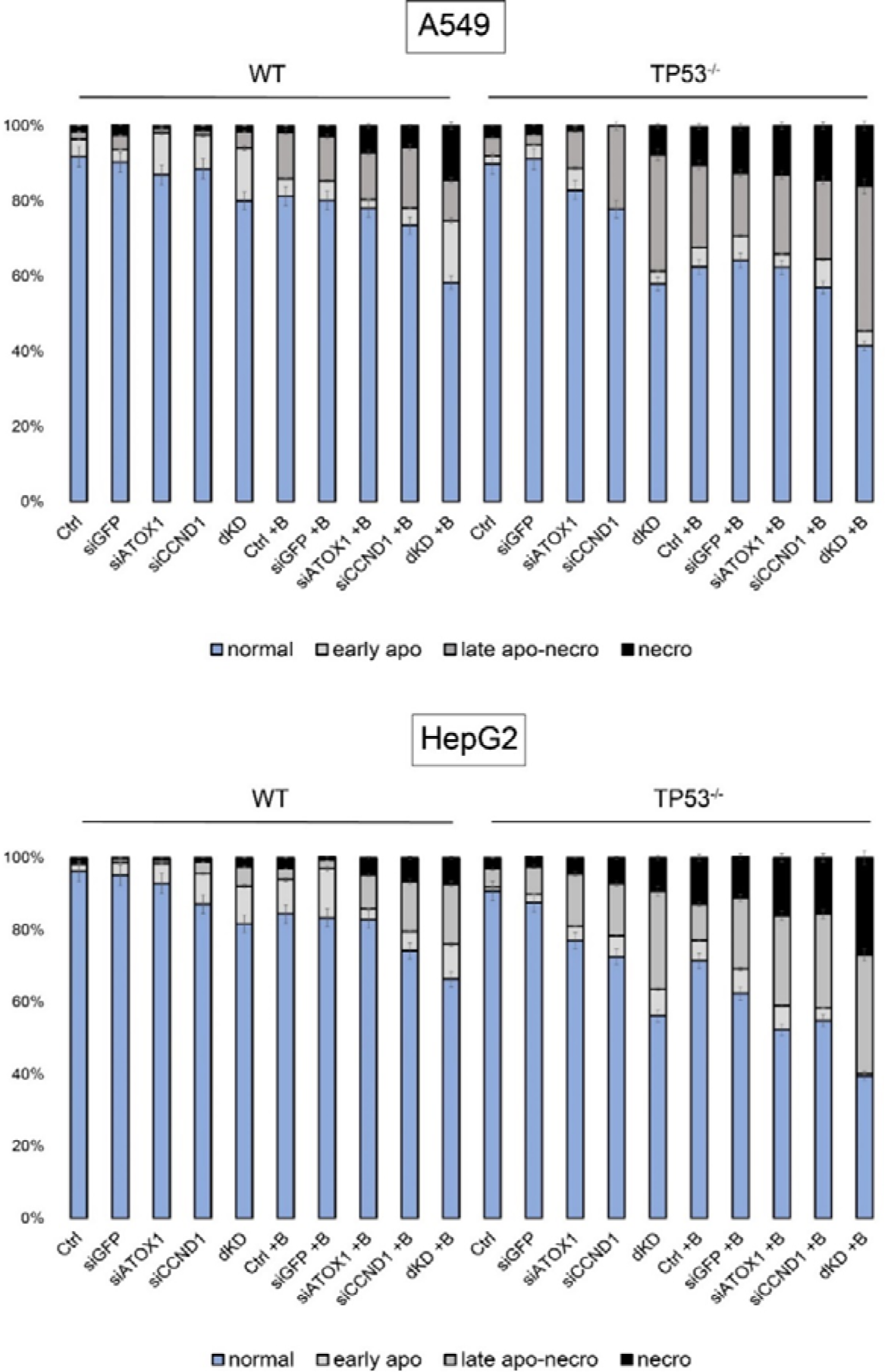
Distribution of cell populations stained with PI and YO-PRO-1 following sequential siRNA inactivation (siCCND1, siATOX1), as well as simultaneous suppression of *CCND1* and *ATOX1* (dKD), under intact conditions and upon addition of the drug Bleo (+B) to A549 and HepG2 cell lines. 72 hours post-treatment with the drug (IC50 for each subline, respectively). Normal - intact cells, early apo - early apoptosis, late apo-necro - late apoptosis and necrosis, necro - necrotic fraction. Mean values from 3 independent measurements ± SEM are indicated, differences are significant (p < 0.05, one-way ANOVA).

For wild-type HepG2 cells, cell death via apoptosis was characteristic for 10-15% of cells treated with siRNA targeting *ATOX1* and *CCND1* individually, which increased to 20% when combined. Treatment with bleomycin enhanced the effect by 10-15%, with the addition of the drug to cells with a double knockout (dKD) resulting in the death of 30% of the population predominantly through an apoptotic pathway. Inactivation of *TP53* in HepG2 cells rendered them more sensitive to the suppression of individual genes and the addition of bleomycin: a double knockout in these cells led to the death of 40% of the cell population, with 10% following a necrotic scenario. The addition of the drug caused the death of 60% of the population, with almost half (∼25-29%) exhibiting necrosis.

Taken together, these results confirm that simultaneous inhibition of *ATOX1, CCND1,* and *TP53* induces a lethal effect, which is potentiated by the action of antitumor compounds. Such an effect directs cells towards predominantly apoptotic cell death with a fraction of necrotic cells, which may be a favorable outcome in the context of therapy for tumors with chemoresistance. Further refinement of effective doses and time points is necessary, along with analysis of Atox1- and cyclin D1-dependent targets to minimize off-target effects and reduce the necrotic response.

## DISCUSSION

This study established the possibility of mutual regulation of three mechanisms that mediate the most important processes in tumor cells: p53, Atox1, cyclin D1. The use of stable (gene deletion using CRISPR-Cas9 technology) and transient (sequence-specific siRNA) inactivation of individual mechanisms made it possible to identify the coregulation of *ATOX1* and *CCND1*/cyclin D1 expression and the role of p53 in their repression. The established molecular connections are important for the main criterion for the effectiveness of antitumor effects - the elimination of tumor cells: combined suppression of *ATOX1* and *CCND1* with non-functioning p53 led to spontaneous death; the combination of three conditions caused a pronounced lethal effect when exposed to palbociclib or bleomycin. From a methodological point of view, it is important to use different methods of inactivating gene expression, as well as cell lines that differ in tissue origin and different pharmacological agents: the effects of gene deletion may differ compared to the action of siRNA, tissue-specific features determine differences in responses to individual external factors. At the same time, the coregulation of *ATOX1* and *CCND1* was reproduced by different methods of *TP53* inactivation in the lung and liver tumor cells, and the responses to chemotherapeutic effects in different experimental models were comparable. Previously, in the A549 and HCT116 lines, we have already shown the dependence of the Atox1 level on the status and activity of p53 and p21 [37]. Experiments with siRNA-mediated inactivation of *TP53* on A549 are not repeated in this study.

The relationships identified at the mRNA level are confirmed by studying the corresponding proteins: suppression of *ATOX1* expression entails a decrease in cyclin D1, and vice versa. These patterns occur in cells with intact p53; dysfunctional p53 does not alter the mutual negative regulation of *ATOX1* and *CCND1*, but in this situation the levels of Atox1 and cyclin D1 proteins increase.

The role of Atox1 as a transcription factor activating *CCND1* was demonstrated by group of Itoh et al. [33]. The influence of Atox1 can be exerted directly and/or indirectly through activation of mitogen-activated protein kinases (MAPK) [46] or nuclear factor kappa B (NFκB) [47]. It is important to highlight the involvement of Atox1 in the assembly and functioning of the APC complex (anaphase control factor of the cell cycle), the activation of which stabilizes CDK4/6 and promotes the transition to S phase. Inhibition of Atox1 disrupts the functioning of APC; the CDK4/6-cyclin D1 complex is degraded, which makes it difficult for cells to exit the G_1_ phase and inhibits further proliferation [48]. The mechanism of cyclin D1 reduction upon Atox1 inhibition remains unknown.

Despite the observed combined regulation that may imply a direct link, the complexity of cell cycle regulation must be considered, suggesting the presence of intermediaries between Atox1 and cyclin D1. Specifically, we hypothesize the involvement of CDK4/6 in the joint regulation of these genes. This assumption is based on cyclin D1 direct or indirect participation in the regulation of factors [49] capable of influencing *ATOX1* expression. In particular, cyclin D1 is directly linked to the activity of Dmp1, a transcription factor that induces Arf/p53-dependent cell cycle arrest in the G_2_/M phase followed by apoptosis, with observed inhibition of the MAPK cascade [50].

The role of the p53-Atox1-CCND1 axis in cell viability has been demonstrated for the first time. In cells with intact p53, inhibition of *ATOX1* and *CCND1* individually causes minimal toxicity. Conversely, double knockdown of these genes increases both spontaneous cell death and cytotoxicity of the anticancer drug bleomycin. The synergistic effect *ATOX1* and *CCND1* knockdowns is significantly greater in the absence of functional p53. Flow cytometry analysis of cell death types predominantly revealed an apoptotic nature, enhanced by bleomycin treatment, but also a small fraction (10-15%) of necrotic cells was observed. The combined influence of Atox1 and cyclin D1 on cell viability has not been previously described, although there are reports on the impact of individual mechanisms. For instance, Atox1 may contribute to cisplatin or doxorubicin resistance by promoting DNA damage repair [51]. An anti-apoptotic role of Atox1 in activating survival signaling mechanisms, particularly MAPK and NFκB has been demonstrated [46, 47]. Inhibition of cyclin D1 is used in therapy for its pro-apoptotic effects [52, 53]. Concurrent suppression of Atox1 and cyclin D1 likely leads not only to cell cycle arrest but also to decreased reparative processes, enhancing the effectiveness of genotoxic drugs. Figure 6 summarizes the studied interactions.

**Fig. 6.**
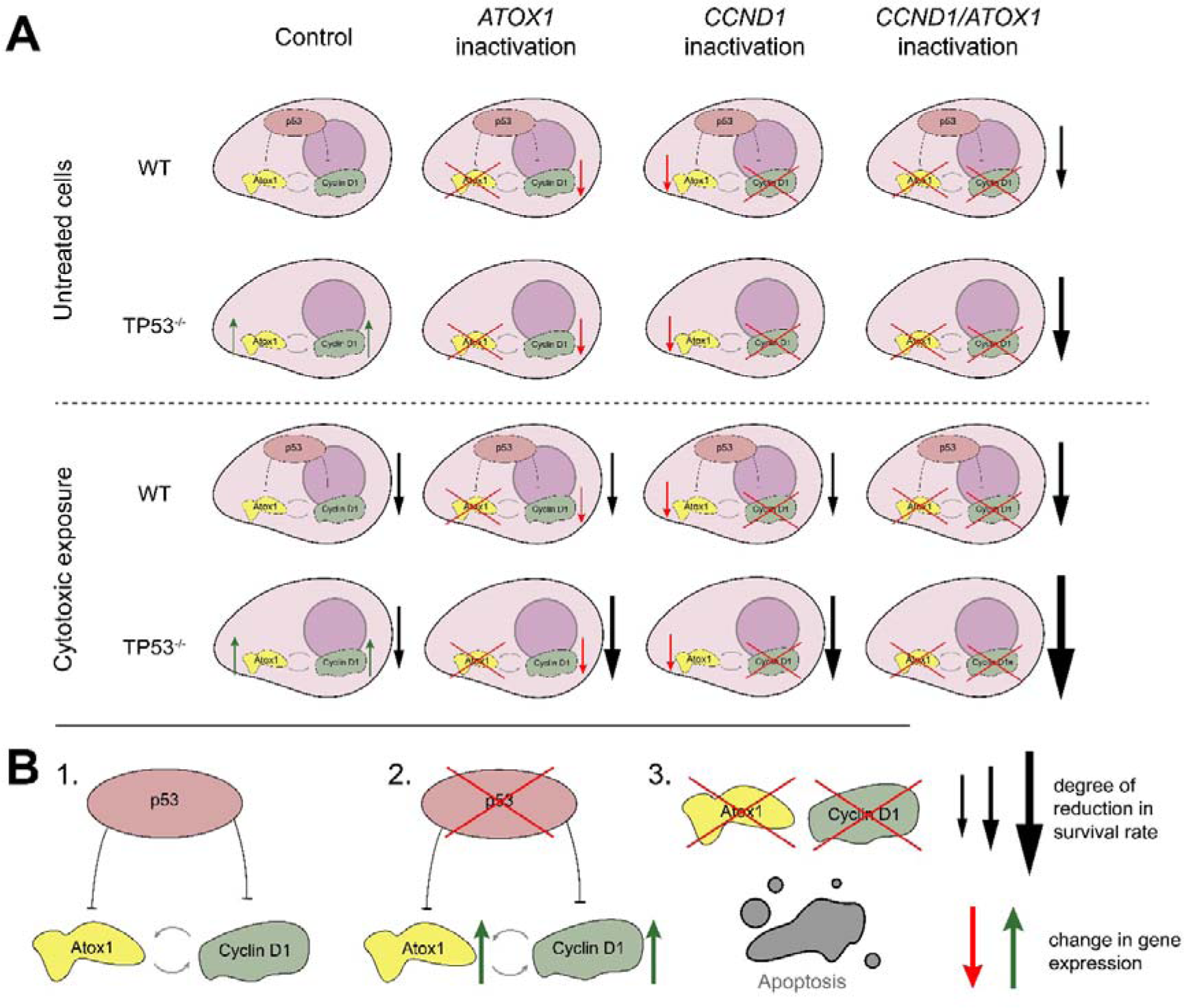
A – diagram of mutual regulation of p53, Atox1 and cyclin D1. B – diagram of the detected interactions between the studied genes and their products: 1. Cyclin D1 and Atox1 are connected by positive feedback, which is masked by active p53. 2. Inactivation of p53 induces cyclin D1 and Atox1. 3. Simultaneous inhibition of *CCND1* and *ATOX1* increases spontaneous death, which is significantly enhanced in response to anticancer drugs. WT – cells with normally functioning p53, TP53^-/-^ – cells with knockout of the *TP53* gene

The identification of gene pairs, the combination of which enhances drug effects – a phenomenon of synthetic lethality [54, 55] – is a crucial area of molecular oncobiology. The results obtained are potentially important for studying the issue of “conditional” (induced by additional interventions) synthetic lethality. Ultimately, knowledge of gene expression profiles in specific tumors is necessary for therapy individualization. These characteristics account for substantial differences in the efficacy of the same drug among individual patients. The negative impact of p53 on the expression of both *ATOX1* and *CCND1* established in this study, as well as the significant enhancement of cytotoxicity when inhibiting these genes in combination with non-functional p53, suggests high effectiveness of targeted suppression of *ATOX1* and CDK4/6-cyclin D1 complexes in tumors with *TP53* mutations. This reveals a new perspective for developing molecular markers for tumor response to treatment. For such situations, the combination of single-target drugs or the development of “dual-action” compounds using PROTAC technology [56] for simultaneous inactivation of *ATOX1* and *CCND1* have been shown. It is important to determine the influence of individual p53 mutations (not only LOF) on the effectiveness of Atox1/Cyclin D1 inhibitors. Identification of the molecular mechanisms of p53-Atox1-Cyclin D1 interaction is required, including direct transcriptional regulation and influences mediated by MAPK and NFκB cascades. Attention should be paid to the contribution of reactive oxygen species (ROS) to Atox1 and cyclin D1 activity, as Atox1 is involved in regulating the redox balance by stimulating ROS production through NADPH oxidase activation. This contradicts the initial hypothesis about copper chaperones as an ROS elimination factors, but explains its p53-dependent suppression [32, 57, 58]. In conclusion, our study highlights the need for a broader description of the relationship between *ATOX1* and copper metabolism proteins with the cell cycle control and repair machinery, which will ultimately lead to the discovery of new therapeutically relevant targets for combined treatment of tumors with various functional oncogene mutations.

## Acknowledgments

The authors express their sincere gratitude to Bruter A.V. (Institute of Gene Biology, Russian Academy of Sciences) for providing the A549TP53^-/-^ cell line and Dr. Shtil A.A. (N.N. Blokhin National Medical Research Center of Oncology) for critical discussion of the manuscript.

## Compliance with ethical standards

This article does not contain any studies with human participants or animals performed by any of the authors.

## Authors’ contributions

Kuchur O.A. and Tsymbal S.A. – concept and management of the work, discussion of the research results, writing and editing the text, working with drawings; Pogodaeva S.S. and Shcherbakova A.V. – conducting experiments, statistical data processing

## Abbreviations

Bleo: bleomycin
Palbo: palbociclib
siRNA: small interfering RNA
siGFP: control siRNA to the *GFP* gene
TP53^-/-^: cells with a knockout of the *TP53* gene
WT: wild-type cells
qPCR: real-time PCR

